# Stem cell memory EBV-specific T cells control post-transplant lymphoproliferative disease and persist *in vivo*

**DOI:** 10.1101/2023.05.30.542809

**Authors:** Darya Palianina, Juliane Mietz, Claudia Stühler, Brice Arnold, Glenn Bantug, Christian Münz, Obinna Chijioke, Nina Khanna

## Abstract

Adoptive T cell therapy (ACT), the therapeutic transfer of defined T cell immunity to patients, offers great potential in the fight against different human diseases including difficult-to-treat viral infections but response rates are still suboptimal. Very early differentiated stem cell memory T cells (T_SCM_) have superior self-renewal, engraftment, persistence, and anti-cancer efficacy, but their potential for anti-viral ACT remains unknown. Here, we developed a clinically-scalable protocol for expanding Epstein-Barr virus (EBV)-specific T_SCM_-enriched T cells with high proportions of CD4^+^ T cells and broad EBV antigen coverage. These cells showed tumor control in a xenograft model of post-transplant lymphoproliferative disorder (PTLD) and were superior to previous ACT protocols in terms of tumor infiltration, *in vivo* proliferation, persistence, proportion of functional CD4^+^ T cells, and diversity of EBV antigen specificity. Thus, our new protocol may pave the way for the next generation of potent unmodified antigen-specific cell therapies for EBV-associated diseases, including tumors, and other indications.

## INTRODUCTION

T cell therapies are promising for treatment of hemato-oncological diseases,^1,2^ difficult- to-treat viral infections, and autoimmune diseases.^3,4^ The efficacy of these therapies depends on T cell activation by antigens and *in-vivo* persistence for sustained impact.^5^ Activated T cells can differentiate to stem cell memory (T_SCM_), central memory (T_CM_), transitional memory (T_TM_) effector memory (T_EM_), and terminally differentiated, short-lived effector T cells (T_EMRA_).^6^ During T cell differentiation, effector functions increase, but self-renewal capacity declines.^7^ The superior proliferation and persistence of T_SCM_ has been demonstrated after adoptive transfer of genetically modified lymphocytes,^8^ chimeric antigen receptor (CAR) T cells, engineered T-cell receptor (TCR)-T cells, and tumour-infiltrating lymphocytes (TILs).^9,10,11^ Long-lasting antigen-specific T_SCM_ were also identified after yellow fever and bacillus Calmette–Guerin (BCG) vaccination.^12,13^ CD8^+^ T_SCM_ support T-cell responses to chronic LCMV infection^14^ and are associated with improved prognosis in chronic HIV-1 infection.^15^

T_SCM_ might also offer exciting avenues to improve adoptive therapy with virus-specific T cells (VST) against viral infections that are important causes of morbidity and mortality of immune-deficient transplant recipients. Adoptive transfer of VST can restore virus-specific immunity and prevent or cure such viral infections.^16,17^ This includes transfer of EBV-specific cytotoxic T-cell lines (CTLs) that prolongs overall survival in patients with EBV-driven post-transplant lymphoproliferative disease (PTLD), other EBV-associated lymphomas and possibly even immunopathologies due to inefficient EBV specific immune control. However, ∼30% of patients show no response indicating a need for further improvements.^18^ Limited long-term response because of poor persistence and exhaustion of the transferred T cells might contribute to insufficient response rates. Most clinical studies used VST generated by long-term expansion with continuous re-stimulation with EBV-antigen expressing lymphoblastoid cell lines (LCLs),^19^ potentially driving the cells to late differentiation stages and exhaustion.^20^ Alternatively, VST can be generated by rapid expansion using a single stimulation with a viral peptide mixture, but the differentiation state and persistence for EBV lymphomas is unknown.^21,22^

Here, we established a novel facile, robust, and clinically applicable protocol for rapid expansion of EBV-CTLs with a high fraction of EBV-specific T_SCM_. These cells mediate EBV control *in vitro* and *in vivo* and are superior to previous VST protocols in terms of tumor infiltration, *in vivo* proliferation, persistence, proportion of functional CD4^+^ T cells, and diversity of EBV antigen specificity.

## RESULTS

### Rapid expansion in presence of IL-4 / IL-7 and TWS-119 yields high T_SCM_ proportions

To generate EBV-CTLs with high proportions of T_SCM_, we modified the rapid expansion approach.^21^ We stimulated PBMC of healthy EBV-seropositive donors with the EBV Consensus peptide pool (Figure 1A). We determined the impact of the cytokines IL-7, IL-15, and IL-21, which promote T cell growth but limit differentiation;^21,23,24^ potassium-rich medium promoting T cell stemness preservation;^25^ and the glycogen synthase kinase-3β (GSK3β) inhibitor TWS119, which induces Wnt-beta-catenin signaling limiting cell differentiation and promoting T_SCM_ generation.^26^ We determined the proportion of T_SCM_ (CD45RA^+^ CD45RO^-^ CD62L^+^ CD27^+^) using flow cytometry (Supplemental Figure 1A).

**Figure 1.**
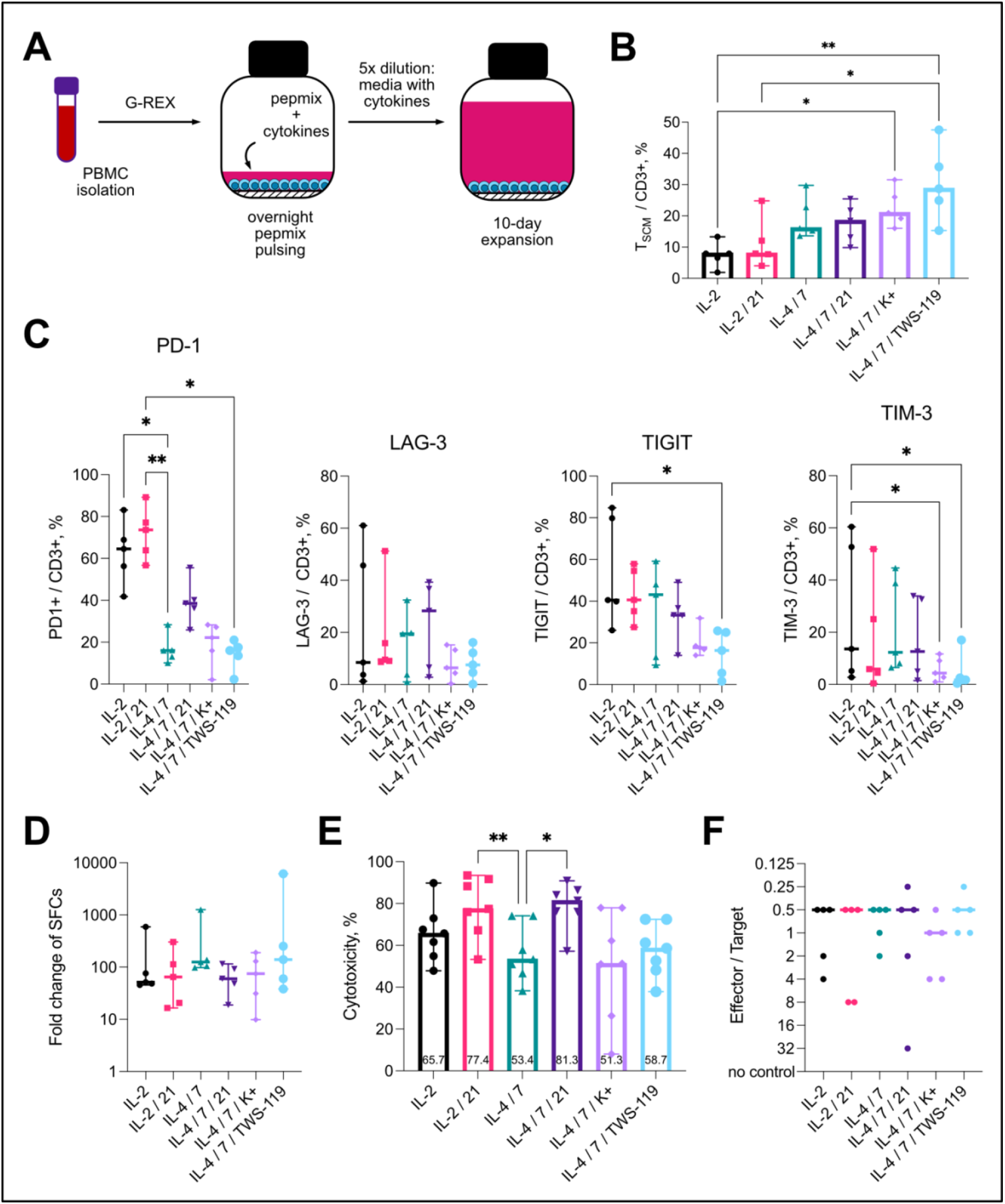
Establishing rapid T_SCM_-enriched EBV-CTL *ex vivo* expansion protocol. (A) Adapted rapid expansion approach. Isolated PBMCs were stimulated with EBV Consensus pepmix in complete media. After overnight pulsing, pepmix was diluted 5x with complete media following 10-day incubation. (B) T_SCM_ proportions after culturing EBV-CTLs in the presence of different conditions (different cytokine combinations, in elevated potassium concentration (K+) or with the edition of TWS-119) as detected by flow cytometry; n=5, medians with range. (C) PD-1, LAG-3, TIGIT and TIM-3, exhaustion marker expression of expanded CTLs; n=5, medians with range. (D) Expansion folds (PBMCs vs. after rapid expansion) of spot-forming cells after culturing in different conditions; IFNγ ELISpot with EBV pepmix stimulation, n=5, medians with range. (E) Short-term cytotoxicity against autologous EBV-LCLs, medians with range. (F) Long-term cytotoxicity: 4-week EBV-LCL outgrowth control by expanded T cells; n=5; medians of controlling E : T were shown. B-F were analyzed by Friedman test, α=0.05, non-significant p-values (ns) not shown, *p < 0.05, **p < 0.005.

IL-4 / IL-7 promoted T_SCM_, whereas IL-21 had limited impact (Figure 1B). Cytokine combinations with IL-15 enriched for NK and NKT cells, and the T_SCM_ proportions were insufficient to consider NK cell depletion for the use in the clinical setting (Supplemental Figure 2A-C). Potassium-rich medium and TWS-119 also promoted T_SCM_. Overall, IL-4 / IL-7 / TWS-119 yielded the highest T_SCM_ proportion (∼30%) with comparable CD4^+^ and CD8^+^ T cell subsets (Supplemental Figure 1B). This condition also showed the lowest levels of T cell exhaustion markers (Figure 1C). The different conditions had limited impact on overall and antigen-specific T cell expansion as measured by total cell counts, ELISpot assays, and MHC class I multimer staining (Figure 1D; Supplemental Figure 3A-C). The optimized protocol yielded EBV-CTLs with reduced short-term cytotoxicity against EBV-lymphoblastoid cell lines (LCLs) (possibly due to delayed activation of early differentiated T cells) but comparable long-term LCL control (Figure 1D-E). Thus, rapid expansion in the presence of IL-4, IL-7 and TWS119 yielded EBV-specific CTLs with favorable properties for virus-specific T-cell therapy including high proportion of T_SCM_, low exhaustion, and efficient long-term *in-vitro* cytotoxicity.

### Expanded T_SCM_ are EBV-specific and proliferate in response to restimulation

To determine the antigen-specificity of various memory T cell subsets within the expanded CTLs, we sorted these subsets (Figure 2A) and co-cultured them with irradiated autologous EBV-transformed LCLs. All memory populations showed proliferation of CD4^+^ and particularly CD8^+^ T cells in these co-cultures (Figure 2B). IFNγ production upon restimulation with EBV peptides confirmed the specificity of proliferating T_SCM_ cells (Figure 2B). This was also consistent with MHC class I multimer staining (Figure 2C).

**Figure 2.**
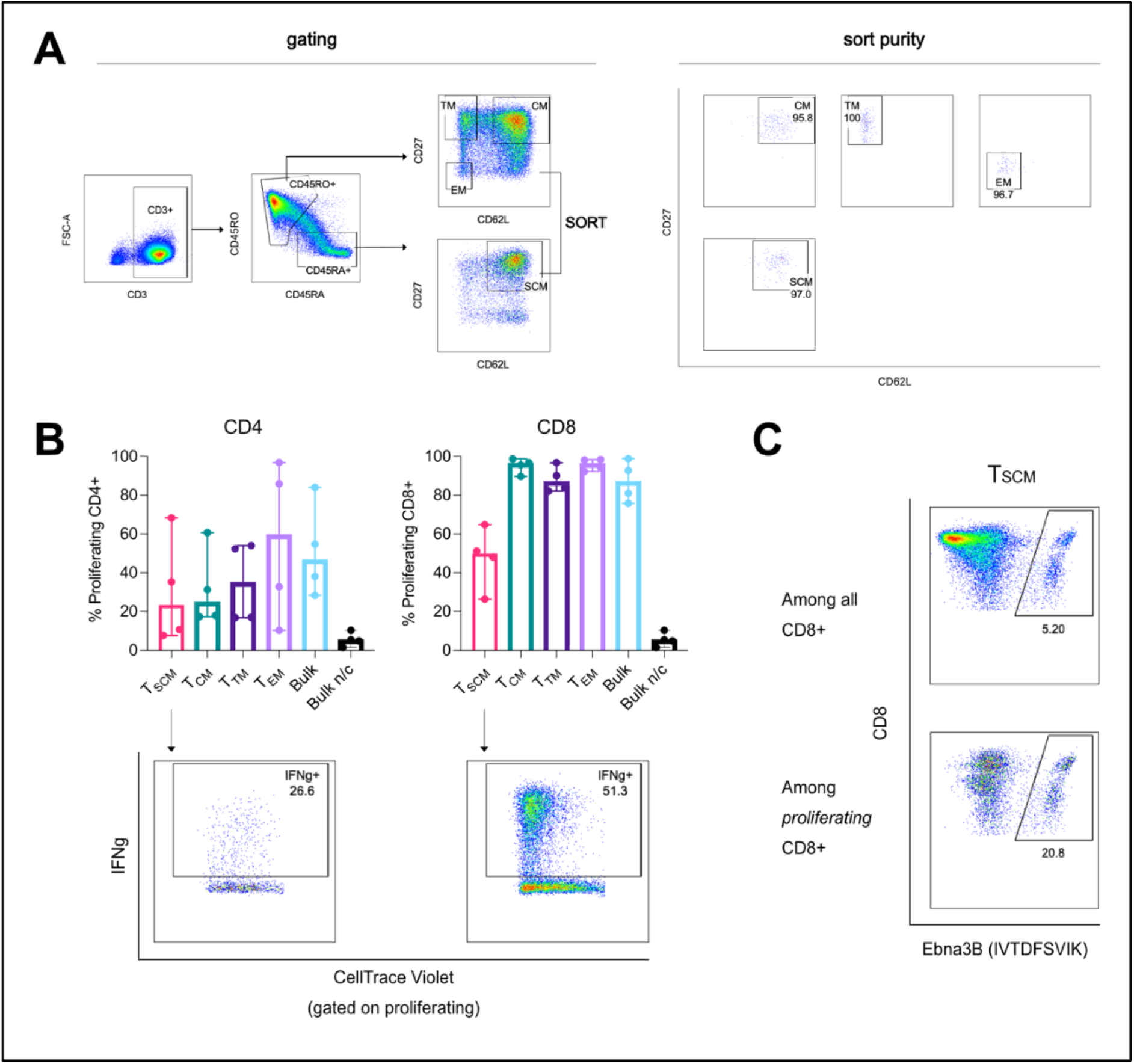
EBV-specific T cells among T_SCM_. (A) Sorting of different memory population: gating strategy and representative plots of sort purity. (B) Proliferation of sorted and CTV-stained CD4^+^ and CD8^+^ T cell populations after 7-day co-culture of the sorted populations with EBV-LCLs (flow cytometry, n=4, medians with range) and specificity of proliferating T_SCM_ cells (IFNγ expression upon re-stimulation with EBV pepmix, representative plot). (C) Proliferation of specific T_SCM_ (stained with a respective MHC class I multimer, representative plot. Proliferating cells in B-C are cells that proliferated at least once or more. SCM – stem cell memory, CM – central memory, TM – transitional memory, EM – effector memory, n/c – no co-culture.

### T_SCM_-enriched EBV-CTLs have a favorable phenotype and broad specificity

The most widely used clinical protocol employs EBV-transformed LCLs as antigen-presenting cells (APCs) for expanding EBV CTLs in 4-5 week-long co-cultures^18,19^ (CTL-L) (Figure 3A). We compared the established protocol and the newly developed T_SCM_-enriching protocol (CTL-R) to evaluate the differences regarding specificity and phenotypes.

**Figure 3.**
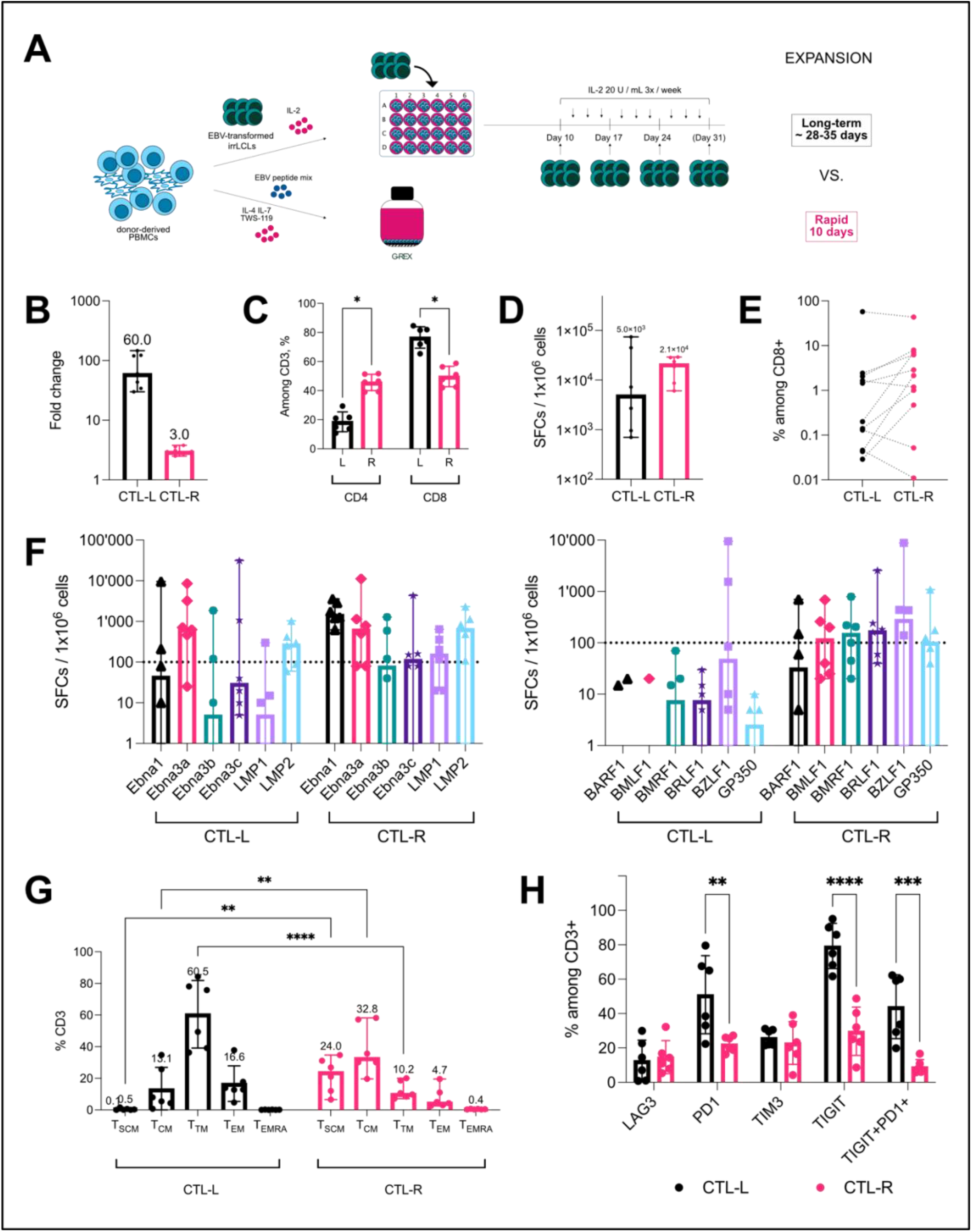
Comparison of rapidly expanded (CTL-R) and long-term conventionally expanded (CTL-L) EBV-CTLs. (A) Schematic of two expansion methods. (B) Expansion rates of total cells. n=7, medians with range. (C) CD4^+^ and CD8^+^ T cell proportions in expanded cells. 2way ANOVA, n=6, means with standard deviation (SD). (D) Frequencies of EBV-specific T cells in the expanded products, IFNγ ELISpot after re-stimulation with EBV pepmix; n=6, medians with range, Wilcoxon matched pairs signed-rank test. (E) Pair-wise comparisons of proportions of different single EBV antigen-specific T cells measured by respective MHC class I-multimer staining, flow cytometry. n=11, Wilcoxon matched pairs signed-rank test. (F) Frequencies of single protein-specific T cells in the expanded products (latent – left graph, lytic – right graph), IFNγ ELISpot after re-stimulation with peptide pools derived from single EBV proteins, the dotted line indicates the threshold (spot calculations below the line were considered not significantly different from control). n=6, medians with range, Wilcoxon matched pairs signed-rank test. (G) Memory phenotypes and (H) exhaustion marker expression, flow cytometry. n=6, means with SD, 2way ANOVA. For C-H: α=0.05, non-significant p-values (ns) not shown, p < 0.05, **p < 0.005, ***p < 0.001, ****p < 0.0001.

CTL-L showed a higher T cell expansion with a lower proportion of CD4^+^ T cells than CTL-R (Figure 3B-C). Overall, EBV specificity was comparable (Figure 3D-E), but CTL-R had broader antigen specificity for both latent and lytic peptides (Figure 3F). Memory phenotypes differed substantially with higher proportions of earlier differentiation stages (T_SCM_, T_CM_) in CTL-R and later differentiation stages (T_TM_, T_EM_) in CTL-L (Figure 3G) and lower levels of exhaustion markers in CTL-R (Figure 3H). Thus, the standard protocol yields more cells, but the novel protocol yields a broader antigen specificity and more favorable memory and exhaustion phenotypes.

### T_SCM_-enriched EBV-CTLs control tumor growth, proliferate, persist and release pro-inflammatory cytokines *in vivo*

To test the *in-vivo* function of T_SCM_-enriched EBV-CTL, we used a well-characterized mouse model of EBV-driven post-transplant lymphoproliferative disease (PTLD).^27,28^ 2x10^6^ luciferase-expressing EBV-LCLs were injected subcutaneously followed by adoptive transfer of 1x10^7^ autologous CTL-L or CTL-R three days later i.v. into NOD-*scid* gamma ^-/-^ (NSG) mice supplemented with high doses of human IL-2 to support T cells in the NSG system (Figure 4A). Tumor growth dynamics revealed that both CTL-L and CTL-R controlled tumor growth equally well over four weeks (Figure 4B-C). Three out of 13 mice (23%) receiving CTL-R but no mice receiving CTL-L lost weight at late time points (supplemental Figure 5) together with increased levels of white blood cells (WBCs) and higher serum IFNγ and TNFα levels in the CTL-R group (Figure 4D,G). Spleen weights and splenocyte counts were also higher for CTL-R than CTL-L and tumor-only groups (Figure 4E, supplemental Figure 4A) and spleen, peripheral blood and bone marrow contained more human CD45^+^ (hCD45^+^) cells. Most of these cells were CD3^+^ indicating substantial *in-vivo* expansion of T cells (Figure 4F, supplemental Figure 4B). CD8^+^ T cells expanded initially more in mice receiving CTL-R but CD8^+^/CD4^+^ T cell ratios returned to pre-infusion levels at later time points, whereas CD8^+^ T cells dominated throughout in mice receiving CTL-L group (Figure 4F, supplemental Figure 4C).

**Figure 4.**
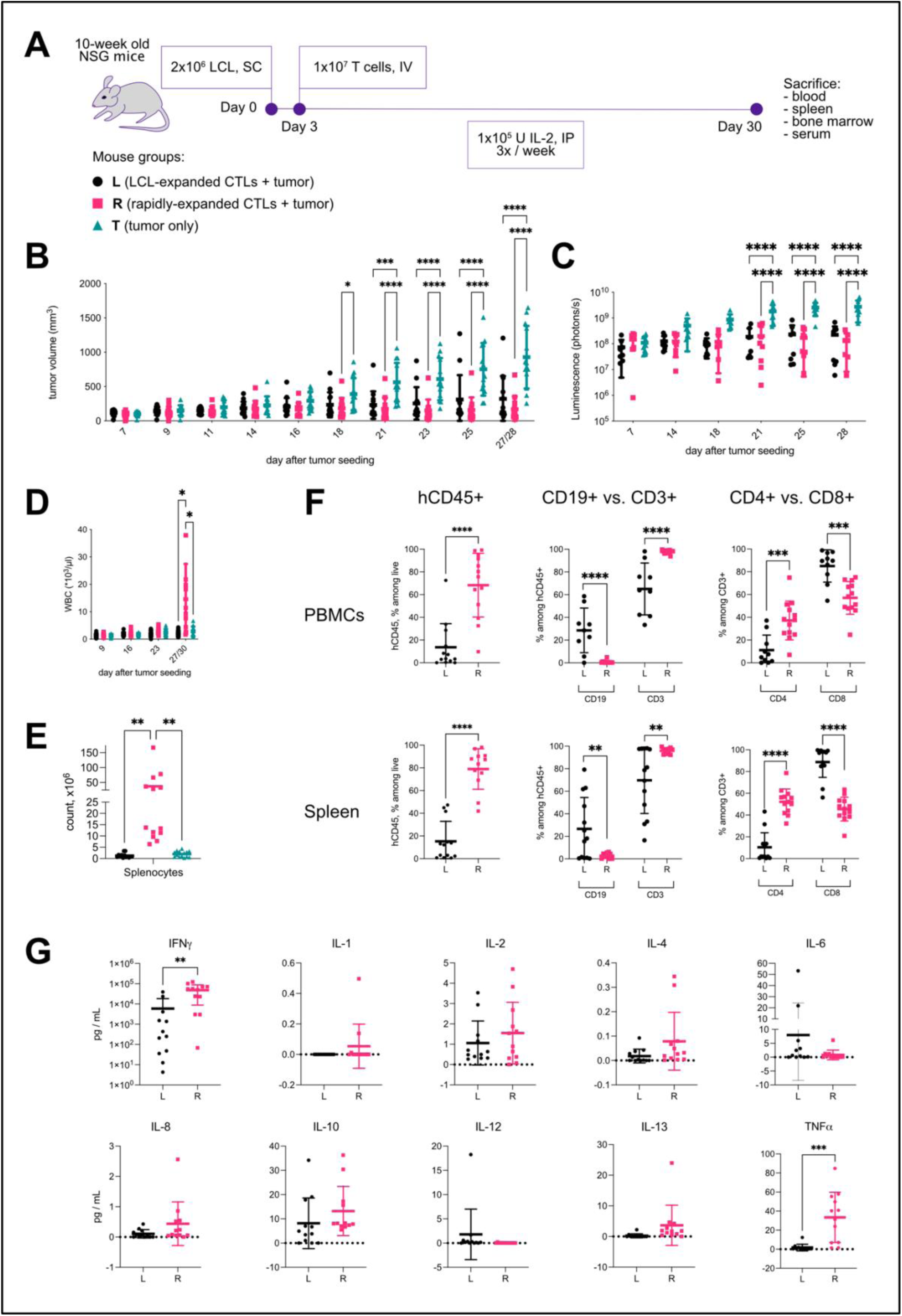
Expansion of CTL-R *in vivo*. (A) Schematic of the *in vivo* experiments. 2x10^6^ tumor cells (luciferase-expressing EBV-LCLs) / mouse were injected into NSG mice subcutaneously, and on day 3 1x10^7^ autologous long-term or rapidly expanded EBV-CTLs per mouse were infused intravenously. Groups of tumor-only mice were kept as a negative control. All mice were supplemented with 1x10^5^ U / hIL-2 3x / week. Mice were sacrificed after ∼4 weeks and organs were collected. Pooled data from three independent experiments with 3 different donors (n = 13 mice/group) is shown in further plots unless there was no sample available. Data from CTL-L-injected mice marked in black circles, CTL-R – in pink squares, and tumor-only mice – in green-blue triangles. Tumor growth dynamic measured by calipering ∼3x / week (B) and tumor luminescence measured 1-2x / week (C). (D) *In vivo* white blood cell (WBC) expansion dynamic, flow cytometry of weekly bleedings. (E) Splenocyte total counts after sacrifice. (F) Proportions of human CD45^+^ cells, CD3^+^ and CD19^+^ among human CD45^+^ cells, and CD4^+^ and CD8^+^ among human CD3^+^ in peripheral blood (PBMC) and spleen, flow cytometry. (G) Multiplex analysis of human cytokines in the murine sera collected after sacrifice. For B-F, means with SD; mixed-effects analysis (B-D, G), 1-way ANOVA and Tukey’s multiple comparrisos test (E) and multiple unpaired t test (F), α=0.05, non-significant p-values not shown, p < 0.05, **p < 0.005, ***p < 0.001, ****p < 0.0001.

Thus, both long-term and rapidly expanded EBV CTLs efficiently control tumor growth, but T_SCM_-enriched CTLs generate more CD4^+^ and CD8^+^ T cells and persist better *in vivo*.

### T_SCM_-enriched EBV CTLs infiltrate tumors and have broad antigen specificity

To determine the specificity of EBV-CTLs expanded *in vivo*, we measured tumor infiltration of CD8^+^ T cells by immunohistochemistry (Figure 5A-B). However, flow cytometry revealed a significantly higher CD3^+^ T cell infiltration and particularly a higher infiltration of CD4^+^ T cells into tumors of mice receiving transfer of CTL-R compared to CTL-L (Figure 5C-D).

**Figure 5.**
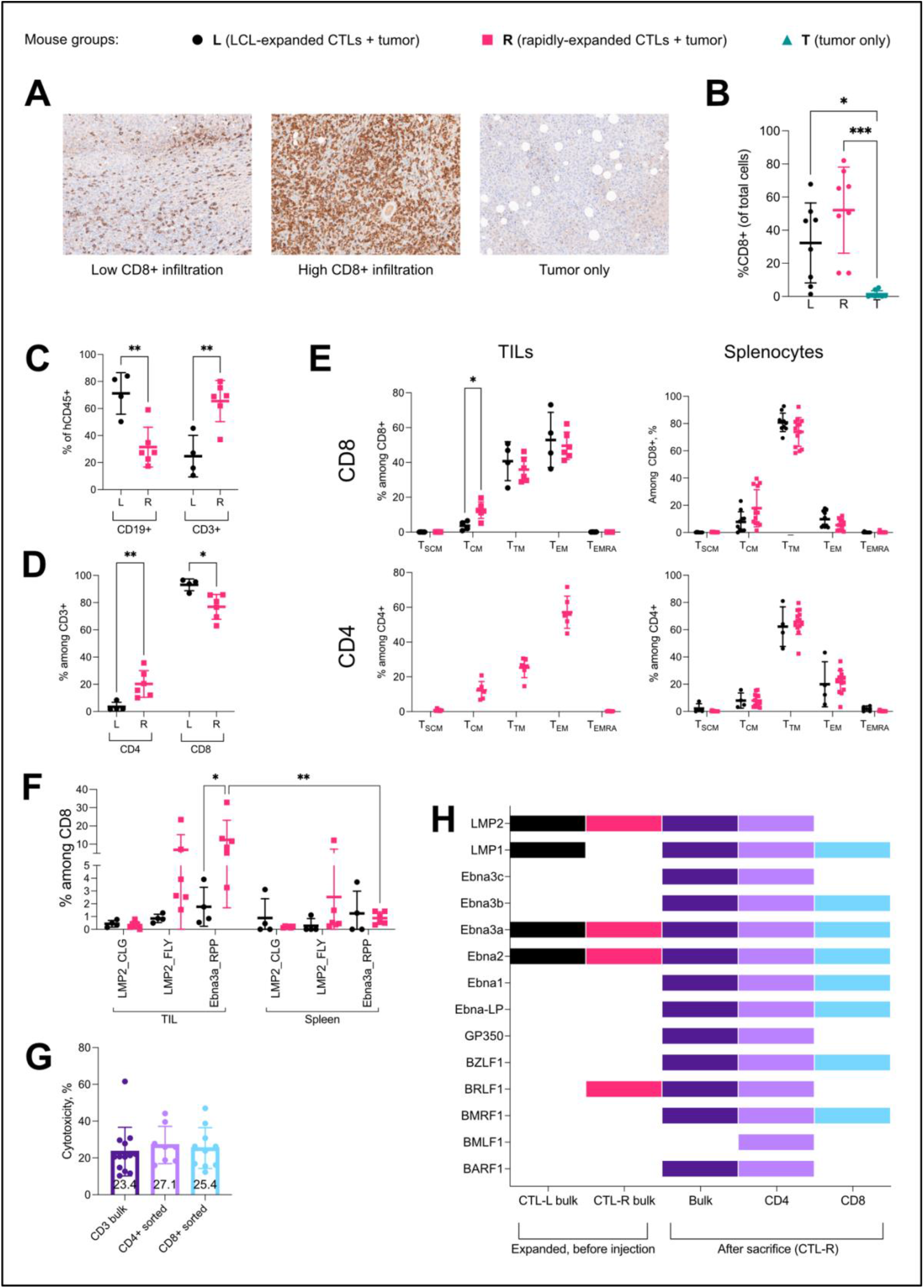
Specificity of expanding EBV-CTLs *in vivo*. Immunohistochemistry analysis of CD8^+^ tumor infiltrating lymphocytes: (A) representative pictures of samples with low and high CD8^+^ T cell infiltration as well as tumor only, CD8^+^ cells are stained in brown, nuclei – in blue; (B) proportions of tumor infiltrating CD8^+^ lymphocytes by treatment group (pooled data from available samples from two experiments, 10 mice / group). (C) Proportions of CD3^+^ vs. CD19^+^ among human CD45^+^ cells in TILs, and (D) proportions of CD4^+^ vs. CD8^+^ among CD3^+^ T cells, measured by flow cytometry. (E) A shift of CD8^+^ and CD4^+^ T cell memory phenotypes in TILs vs. spleen (pooled data of available samples). (F) Proportions of EBV-specific MHC class I-multimer-stained CD8^+^ T cells. (G) *in vitro* short-term cytotoxicity of bulk, separated CD4^+^ and CD8^+^ splenocytes against autologous LCLs (pooled data from all available samples). (H) Presence of specific response of bulk CTL-L and CTL-R cultures before injection vs CTL-R bulk, CD4^+^ and CD8^+^ splenocytes after sacrifice to stimulation with single-EBV protein antigen pools measured by ELISpot (based on mean data from all available samples collected from one experiment / one donor). For B-G, means with SD; 1-way ANOVA and Tukey’s multiple comparrisos test (B, G); multiple unpaired t test (C-E) and 2way ANOVA (F) were used.

CTL-R-derived tumor-infiltrating lymphocytes (TILs) showed a less differentiated phenotype with more T_CM_ compared to CTL-L (Figure 5E), although T_SCM_ TILs were rare in both groups. Tumor-infiltrating T cells of both groups were more differentiated with a predominance of T_EM_ compared to T cells not derived from tumors, with high proportions of T_TM_ in spleen and other organs (Figure 5E, Supplemental Figure 6A-C). A higher proportion of CTL-R-derived antigen-specific TILs was detected via specific EBV peptide loaded MHC class I multimers compared to the CTL-L group and CD8^+^ T cells in spleen from both groups (Figure 5F).

Both CD4^+^ and CD8^+^ splenocytes derived from CTL-R showed short-term *in-vitro* cytotoxicity against EBV-LCLs (there was insufficient expansion in the CTL-L group to perform similar experiments) (Figure 5G). ELISpot assays with peptides from 14 different EBV proteins revealed a remarkable increase of antigen coverage in CTL-R derived splenocytes and particularly CD4^+^ T cells (Figure 5H). Thus, transferred EBV-specific T_SCM_-enriched CTLs (CTL-R) showed robust proliferation and longevity, and reconstituted a wide antigen diversity of different T cell compartments.

## DISCUSSION

Adoptive T cell therapy (ACT) therapeutically transfers specifc T cell immunity to patients. Persistence of memory T cell subsets in the recipients is often critical for long-term efficacy, but difficult to attain. Very early differentiated T_SCM_ cells, characterized by high self-renewal, engraftment and persistence, can reconstitute all types of effector and memory T cell subsets^29^ and show encouraging results in T cell therapy.^8,10,11^

T_SCM_ might be also a promising avenue for treatment of viral infections such as EBV in transplant patients. Anti-viral ACT is an active field with several ongoing phase III trials (clinicaltrials.gov: NCT03394365, NCT04832607, NCT04832607, NCT04832607). However, current strategies are still suboptimal with response rates of only around 70% for most viruses.^18^ To test T_SCM_ for ACT against EBV infections, T cell priming and several steps of cell sorting can be used, but the complexity of this method makes it difficult to translate to the clinical setting.^30^ Moreover, CD4^+^ T cells are depleted which might impair sustaining adoptive immunity.^31^

Here, we developed a novel method to enrich both CD4^+^ and CD8^+^ T_SCM_ EBV-CTL from PBMCs with minimal cell handling steps. We did not use the common stimulation with autologous EBV-LCLs,^32^ which requires 4 to 8 weeks and yields predominantly late-stage T_EM_.^33,34^ Continuous re-stimulation of T cells can promote T cell exhaustion^20^ defined as a reduced functional capacity.^35^ Instead, we adapted a rapid expansion protocol that is safe and effective in transplant patients^36^ and yields a higher proportion of T_CM_^21^, which are superior to T_EM_ in antiviral activity and persistence.^37,38^ Due to its minimal cell handling steps, this protocol can be easily transferred to the clinic. The overall yield is lower than for long-term stimulation, but this is not a limiting factor because sufficient starting numbers of PBMCs can be obtained with standard blood donations.

We systematically tested various cytokines and other conditions to maximize the proportion of T_SCM_ and discovered that a combination of IL-4 and IL-7 with induction of the Wnt/β-catenin pathway using TWS119 triggered efficient enrichment or EBV-specific T_SCM_ with broad coverage of antigens including lytic antigens. T cells specific for lytic antigens can be relevant for treatment of diseases like nasopharyngeal carcinoma (NPC) in a therapeutic setting^39, 40^ as well as prophylaxis of EBV-associated B cell malignancies and NPC, as the lytic phase of EBV contributes to oncogenesis.^41-43^ T cells specific for lytic antigens are difficult to obtain with long-term stimulation using EBV-LCLs which exhibit latency III and express predominantly latent EBV antigens.^44^ A wide diversity of antigen specificity also broadens the scope of cell therapy and reduces the risk of relapse due to antigen escape.^45^ Moreover, lytic EBV antigen-specific CD8^+^ T cells require early memory differentiation with maintained CD27 expression for their protective function,^46^ and this is ensured by our rapid expansion protocol.

A key aspect of an ACT product is the balance between CD4^+^ and CD8^+^ T cell populations,^47,48^ and a high CD4^+^ T cell proportion is associated with better responses to anti-EBV ACT for treatment of PTLD.^49^ In contrast to the widely used long-term stimulation with low yields of CD4^+^ T cells, our novel protocol enriched CD4^+^ and CD8^+^ EBV-specific T cells in all memory populations. The high CD4^+^ T cell proportion was maintained after adoptive transfer and was reflected in tumor infiltration, and CD4^+^ T cells recovered a broad antigen-specific profile *in vivo*. These EBV-specific CD4^+^ T cells may contribute to control of various EBV diseases.

Limitations of this study include lack of long-term data beyond 4 weeks in mice. High human T-cell engraftment in murine organs may lead to xeno-GVHD,^50^ signs of which (e.g., weight loss) were observed in some mice in the CTL-R group. Nevertheless, the efficient CTL-R infiltration into tumors, the *in vitro* cytotoxicity and the specific responses of splenocytes indicated high specificity of the expanded T cells. Moreover, this mouse model-related aspect may not be indicative of potential issues in a clinical application because rapidly expanded virus-specific T cells are safe upon the adoptive transfer into patients.^22^

In conclusion, we demonstrate that our novel protocol yields promising EBV-specific T_SCM_-enriched CTLs with favorable properties for VST production, such as early differentiated memory composition, low exhaustion, high tumor infiltration, efficient CD4^+^ and CD8^+^ T cell mediated cytotoxicity, long-term persistence, and broad antigen specificity. This may pave the way for the next generation of unmodified antigen-specific cell therapies against viral infections. The safety and efficacy as well as the clonal diversity of these VST remain to be investigated in an upcoming clinical trial.

## METHODS

### Peptides

The PepTivator EBV Consensus peptide pool (Miltenyi Biotec), and single peptide pools from various EBV antigens (latent: EBNA-LP, EBNA2, EBNA3a, EBNA3b, EBNA3c, LMP1; lytic: BARF1, BMLF1, BMRF1, BRLF1, BZLF1, GP350/GP340) (JPT Peptide Technologies) were used for T cell stimulation.

### Blood donors, cell culture and generation and expansion of EBV-specific T-cell lines

Blood was obtained after informed consent from healthy donors in accordance with the Declaration of Helsinki. The study was approved by the local ethic committee (Ethikkommission Nordwest-und Zentralschweiz, Project ID PB_2018-00081). Donors were typed for HLA class I and class II alleles. Human peripheral blood mononuclear cells (PBMCs) were isolated from EDTA blood of healthy donors and ^51^ EBV-transformed lymphoblastoid cell lines (LCL) were generated and cultured in LCM-10 media according to previously published protocols^52^ (Supplemental Materials). Long-term EBV-CTL expansion with LCL re-stimulations and rapid expansion protocols were adapted from previously described protocols.^19 21^ All T cells were expanded in CTL-M (Supplemental methods). For rapid expansion, PBMCs were cultured in a G-Rex bioreactor (Wilson&Wolf). 3x10^6^ PBMCs / well of a 24-well G-Rex plate or 1.5x10^7^ / well of a 6-well G-Rex plate were cultured. On day 0, cells were pulsed overnight in CTL-M (or CTL-M with high K^+^ when applicable) containing the EBV Consensus peptide pool (pepmix) and supplemented with cytokines (and TWS-119 when applicable). Afterwards the pepmix (and TWS-119 if applicable) was diluted 5x with CTL-M supplemented only with cytokines. Cell culture went on up to day 10-12 without further supplementation.

For long-term EBV-CTL expansion, PBMCs were stimulated with autologous EBV-LCLs at effector : target (E:T) = 40:1 for 10 days (2x10^6^ PBMCs/well of a 24-well cell culture plate) without cytokine supplementation. Afterwards T cells were re-stimulated weekly at E:T=4:1) and supplemented with 20 U/mL IL-2 3x / week until day 28-35.^19^ LCL culture is described in Supplemental Methods.

### EBV-LCL generation and culture

PBMCs were incubated with recombinant B95-8 or B95-8-fLuc EBV strains (both gifts from Dr. Wolfgang Hammerschmidt, Helmholtz Center Munich, Germany), cultured in LCM-10 media and were treated with 2 µg/ml Cyclosporin A (Sigma Aldrich) and 2 µg/ml CpG ODN 2006 (InvivoGen) weekly until the transformation. Non-irradiated LCLs were always cultured in LCM-10 media (including cytotoxicity and outgrowth assays).

### Co-culture with autologous EBV-LCLs

After fluorescence-assisted cell sorting (FACS) (staining described below), sorted cells were recovered for 3 days in CTL-M supplemented with IL-4 and IL-7. Afterwards, autologous LCLs were irradiated and T cells were stained with CellTrace Violet (CTV). Irradiated LCLs were cultured with T cells at a ratio 1:1 for one week. Then cells were harvested an analyzed by flow cytometry (see below).

### Short-term and long-term in-vitro cytotoxicity

Short-term 6-hour killing assay and long-term 4-week outgrowth assay were adopted as previously published^53^. Briefly, for killing assay, EBV-CTLs were incubated with target EBV-LCLs at an effector (E) to target (T) ratio = 30:1 for 6 h. Afterwards, cells were stained for viability (Zombie Aqua), apoptosis (CellEvent Blue), CD3 and CD19 surface markers (see the panels below). Cytotoxicity was calculated according to the following formula: 100 – ([V_test_ / V_control_] * 100) where V = % viable (CellEvent^-^ Zombie Aqua^-^) CD19^+^ cells.

For outgrowth assay (long-term cytotoxicity assay), T cells were incubated with EBV-LCLs at different effector / target ratios in triplicates for 4 weeks. The readout was the lowest E/T ratio controlling the outgrowth of LCLs which was determined microscopically and confirmed by flow cytometry.

### IFNγ ELISpot, intracellular cytokine staining and V-PLEX

EBV-responsive T cells were identified by stimulation with EBV peptides. Enzyme-linked immunospot assay (ELISpot)^53^ and intracellular cytokine (ICC) staining for flow cytometry detection^54^ were done as previously published.

Human cytokine presence in murine blood sera was analyzed using V-PLEX human pro-inflammatory panel-1 and detected by Mesoscale system according to manufacturer’s instructions.

### Immunomagnetic cell sorting

CD4^+^ and CD8^+^ T cells were isolated using the MACS CD4^+^ / CD8^+^ isolation kit (Miltenyi Biotec) according to the manufacturer’s instructions.

### Immunohistochemistry

Tumors were fixed in a 4% paraformaldehyde solution; further sample preparation and immunohistochemistry staining were done commercially by the Pathology Department of the University Hospital of Basel. Slides were acquired on an automated slide scanning brightfield microscope (Vectra) and positive cells were quantified using inForm automated image analysis software (Akoya Biosciences).

### Flow cytometry and FACS-based cell sorting

All flow cytometry panels are described in detail in the supplemental materials.

If applicable, red blood cells were lysed using ACK (Ammonium-Chloride-Potassium) lysis buffer until the pellet appeared no longer red. If applicable, whole-cell staining for proliferation tracing and viability staining were performed in PBS according to manufacturer’s instructions. Surface staining with antibodies and MHC class I-multimers (if applicable) was performed in FACS buffer (5% FBS, 0.1% NaN3 in PBS).

For intracellular staining, cells were fixed with fixation buffer (Biolegend, 420801) and stained for intracellular markers in the permeabilization buffer (Biolegend, 421002) according to manufacturer’s instructions. For combined intracellular/intranuclear staining, cells were fixed and permeabilized using Transcription-Factor Buffer Set (BD, #562574) according to the manufacturer’s instructions.

Spectral flow cytometry was performed on Cytek Aurora. Fluorescence-assisted cell sorting was performed with BD FACSMelody. Weekly bleedings of mice were analysed with BD LSRFortessa. Data were analyzed using FlowJo software. FlowSOM algorithm was used to define memory T cell populations: stem cell memory (T_SCM_) as CD45RA^+^CD45RO^-^CD62L^+^CD27^+^, central memory (T_CM_) as CD45RA^-^

CD45RO^+^CD62L^+^CD27^+^, transitional memory T_TM_ as CD45RA^-^CD45RO^+^CD62L^-^ CD27^+^, effector memory T_EM_ as CD45RA^-^CD45RO^+^CD62L^-^CD27^-^, and terminally differentiated T_EMRA_ as CD45RA^+^CD45RO^-^CD62L^-^CD27^-^.

### Procedures in vivo

Animal experiments were conducted according to the licence approved by the veterinary office of the canton of Zurich, Switzerland (ZH049/20). NSG (NOD.Cg-*Prkdc*^*scid*^ *Il2rg*^*tm1Wjl*^*/*SzJ (#005557)) or NSG-A2 (NOD.Cg-*Mcph1*^*Tg(HLA-A2*.*1)1Enge*^ *Prkdc*^*scid*^ *Il2rg*^*tm1Wjl*^*/*SzJ (#009617)) mice were purchased from The Jackson Laboratory and bred and housed under specific pathogen-free conditions at the Laboratory Animal Services Center (LASC) of the University of Zurich. Experiments were initiated at 6-12 weeks of age. The mouse models were adapted from previous studies ^27,28^. LCL tumor cells were injected subcutaneously into the left flank under isoflurane narcosis. 2x10^6^ tumor cells were resuspended in PBS and right before injection mixed in a 1:1 V/V ratio with Corning® Matrigel® Growth Factor Reduced (GFR) Basement Membrane Matrix. Three days after tumor injection, 1x10^7^ T cells were adoptively transferred by tail vein injection. T cell expansion was supported by i.p. injection of 10^5^ IU recombinant human IL-2 (3x/week, Peprotech), or as stated otherwise. Tumor size was monitored by calipering (3x/week) and bioluminescent imaging for tumor cells transformed with a luciferase encoding recombinant EBV strain (generous gift of Dr. Wolfgang Hammerschmidt, Helmholtz Institute Munich, Germany; 2x/week). General health was monitored by weighing and health parameter scoring 3x/week or daily, according to the animal license. Peripheral blood composition and expansion of adoptively transferred T cells were monitored by weekly tail vein bleeding and flow cytometric analysis (Supplementary Materials) on BD Fortessa. White blood cell counts were determined from full blood with an automatic cell counter (DxH 500, Beckman Coulter). For bioluminescent imaging, mice were injected with 5µl/g body weight of 15mg/ml VivoGlo™ Luciferin (Promega) and imaged 10 minutes after injection in an IVIS machine (PerkinElmer) under isoflurane narcosis. Animals were euthanized when they met pre-defined criteria stated in the animal license, or when the control group met the end-point criteria.

### Statistics

Analyses were conducted using Prism software (GraphPad). Data of individual donors are shown as representative experiments or medians with standard deviations (SD). Combined data of different donors are given as median with range.

## Data Sharing Statement

For original data, please contact.

## Acknowledgements

We would like to thank the FACS Core facility of the Department of Biomedicine, University Hospital of Basel, FACS Core facility and Animal Facility of the Institute of Experimental Immunology, University of Zurich, for excellent support. We thank Joëlle Handschin for the assistance in the multiplex assay and Prof. Dr. Dirk Bumann for the valuable scientific and writing advice. Big thanks go to all the blood donors participated in the study. This work was supported by Cancer Research Switzerland Grant No. KFS-4371-02-2018 and KFS-5292-02-2021 (to O.C.), the Swiss National Foundation Grant 32003B_204944 (to N.K.), NCCR Antiresist Grant No. 180541, Switzerland (to N.K.), Bangerter–Rhyner Stiftung (to N.K.).

## Authorship contributions

DP, CS and NK designed the study. DP, CS and BA performed *in vitro* experiments and analysis; GB assisted with the EBV-LCL transformation. DP, JM, OC, CM and NK designed *in vivo* experiments; OC and CM supervised *in vivo* experiments; DP and JM performed *in vivo* experiments and analysis; DP and NK wrote the manuscript.

## REFERENCES

1. Maus MV, Fraietta JA, Levine BL, Kalos M, Zhao Y, June CH. Adoptive immunotherapy for cancer or viruses. Annu Rev Immunol. 2014;32:189–225.

2. Rosenberg SA, Restifo NP. Adoptive cell transfer as personalized immunotherapy for human cancer. Science. 2015;348(6230):62–68.

3. Barrett AJ, Prockop S, Bollard CM. Virus-Specific T Cells: Broadening Applicability. Biol Blood Marrow Transplant. 2018;24(1):13–18.

4. Scott DW. Genetic Engineering of T Cells for Immune Tolerance. Mol Ther Methods Clin Dev. 2020;16:103–107.

5. Redeker A, Arens R. Improving Adoptive T Cell Therapy: The Particular Role of T Cell Costimulation, Cytokines, and Post-Transfer Vaccination. Front Immunol. 2016;7:345.

6. Mahnke YD, Brodie TM, Sallusto F, Roederer M, Lugli E. The who’s who of T-cell differentiation: human memory T-cell subsets. Eur J Immunol. 2013;43(11):2797–2809.

7. Joshi NS, Kaech SM. Effector CD8 T cell development: a balancing act between memory cell potential and terminal differentiation. J Immunol. 2008;180(3):1309–1315.

8. Oliveira G, Ruggiero E, Stanghellini MT, et al. Tracking genetically engineered lymphocytes long-term reveals the dynamics of T cell immunological memory. Sci Transl Med. 2015;7(317):317ra198.

9. Wang F, Cheng F, Zheng F. Stem cell like memory T cells: A new paradigm in cancer immunotherapy. Clin Immunol. 2022;241:109078.

10. Fraietta JA, Lacey SF, Orlando EJ, et al. Determinants of response and resistance to CD19 chimeric antigen receptor (CAR) T cell therapy of chronic lymphocytic leukemia. Nat Med. 2018;24(5):563–571.

11. Biasco L, Izotova N, Rivat C, et al. Clonal expansion of T memory stem cells determines early anti-leukemic responses and long-term CAR T cell persistence in patients. Nat Cancer. 2021;2(6):629–642.

12. Fuertes Marraco SA, Soneson C, Cagnon L, et al. Long-lasting stem cell-like memory CD8+ T cells with a naive-like profile upon yellow fever vaccination. Sci Transl Med. 2015;7(282):282ra248.

13. Mpande CAM, Dintwe OB, Musvosvi M, et al. Functional, Antigen-Specific Stem Cell Memory (TSCM) CD4(+) T Cells Are Induced by Human Mycobacterium tuberculosis Infection. Front Immunol. 2018;9:324.

14. Utzschneider DT, Charmoy M, Chennupati V, et al. T Cell Factor 1-Expressing Memory-like CD8(+) T Cells Sustain the Immune Response to Chronic Viral Infections. Immunity. 2016;45(2):415–427.

15. Ribeiro SP, Milush JM, Cunha-Neto E, et al. The CD8(+) memory stem T cell (T(SCM)) subset is associated with improved prognosis in chronic HIV-1 infection. J Virol. 2014;88(23):13836–13844.

16. Houghtelin A, Bollard CM. Virus-Specific T Cells for the Immunocompromised Patient. Front Immunol. 2017;8:1272.

17. Heslop HE, Sharma S, Rooney CM. Adoptive T-Cell Therapy for Epstein-Barr Virus-Related Lymphomas. J Clin Oncol. 2021;39(5):514–524.

18. Walti CS, Stuehler C, Palianina D, Khanna N. Immunocompromised host section: Adoptive T-cell therapy for dsDNA viruses in allogeneic hematopoietic cell transplant recipients. Curr Opin Infect Dis. 2022;35(4):302–311.

19. Rooney CM, Smith CA, Ng CY, et al. Infusion of cytotoxic T cells for the prevention and treatment of Epstein-Barr virus-induced lymphoma in allogeneic transplant recipients. Blood. 1998;92(5):1549–1555.

20. Zou D, Dai Y, Zhang X, et al. T cell exhaustion is associated with antigen abundance and promotes transplant acceptance. Am J Transplant. 2020;20(9):2540–2550.

21. Gerdemann U, Keirnan JM, Katari UL, et al. Rapidly generated multivirusspecific cytotoxic T lymphocytes for the prophylaxis and treatment of viral infections. Mol Ther. 2012;20(8):1622–1632.

22. Gerdemann U, Katari UL, Papadopoulou A, et al. Safety and clinical efficacy of rapidly-generated trivirus-directed T cells as treatment for adenovirus, EBV, and CMV infections after allogeneic hematopoietic stem cell transplant. Mol Ther. 2013;21(11):2113–2121.

23. Pethe K, Sequeira PC, Agarwalla S, et al. A chemical genetic screen in Mycobacterium tuberculosis identifies carbon-source-dependent growth inhibitors devoid of in vivo efficacy. Nat Commun. 2010;1(57):57.

24. Khalaf WS, Garg M, Mohamed YS, Stover CM, Browning MJ. In vitro Generation of Cytotoxic T Cells With Potential for Adoptive Tumor Immunotherapy of Multiple Myeloma. Front Immunol. 2019;10:1792.

25. Vodnala SK, Eil R, Kishton RJ, et al. T cell stemness and dysfunction in tumors are triggered by a common mechanism. Science. 2019;363(6434).

26. Gattinoni L, Zhong XS, Palmer DC, et al. Wnt signaling arrests effector T cell differentiation and generates CD8+ memory stem cells. Nat Med. 2009;15(7):808–813.

27. Koehne G, Doubrovin M, Doubrovina E, et al. Serial in vivo imaging of the targeted migration of human HSV-TK-transduced antigen-specific lymphocytes. Nat Biotechnol. 2003;21(4):405–413.

28. Hiwarkar P, Qasim W, Ricciardelli I, et al. Cord blood T cells mediate enhanced antitumor effects compared with adult peripheral blood T cells. Blood. 2015;126(26):2882–2891.

29. Gattinoni L, Restifo NP. Moving T memory stem cells to the clinic. Blood. 2013;121(4):567–568.

30. Kondo T, Imura Y, Chikuma S, et al. Generation and application of human induced-stem cell memory T cells for adoptive immunotherapy. Cancer Sci. 2018;109(7):2130–2140.

31. Li K, Donaldson B, Young V, et al. Adoptive cell therapy with CD4(+) T helper 1 cells and CD8(+) cytotoxic T cells enhances complete rejection of an established tumour, leading to generation of endogenous memory responses to non-targeted tumour epitopes. Clin Transl Immunology. 2017;6(10):e160.

32. Prockop S, Doubrovina E, Suser S, et al. Off-the-shelf EBV-specific T cell immunotherapy for rituximab-refractory EBV-associated lymphoma following transplant. J Clin Invest. 2019.

33. Ricciardelli I, Blundell MP, Brewin J, Thrasher A, Pule M, Amrolia PJ. Towards gene therapy for EBV-associated posttransplant lymphoma with genetically modified EBV-specific cytotoxic T cells. Blood. 2014;124(16):2514–2522.

34. Heslop HE, Slobod KS, Pule MA, et al. Long-term outcome of EBV-specific T-cell infusions to prevent or treat EBV-related lymphoproliferative disease in transplant recipients. Blood. 2010;115(5):925–935.

35. Wherry EJ, Kurachi M. Molecular and cellular insights into T cell exhaustion. Nat Rev Immunol. 2015;15(8):486–499.

36. Papadopoulou A, Gerdemann U, Katari UL, et al. Activity of broad-spectrum T cells as treatment for AdV, EBV, CMV, BKV, and HHV6 infections after HSCT. Sci Transl Med. 2014;6(242):242ra283.

37. Wang X, Berger C, Wong CW, Forman SJ, Riddell SR, Jensen MC. Engraftment of human central memory-derived effector CD8+ T cells in immunodeficient mice. Blood. 2011;117(6):1888–1898.

38. Klebanoff CA, Gattinoni L, Torabi-Parizi P, et al. Central memory self/tumor-reactive CD8+ T cells confer superior antitumor immunity compared with effector memory T cells. Proc Natl Acad Sci U S A. 2005;102(27):9571–9576.

39. Chung YL, Wu ML. Clonal dynamics of tumor-infiltrating T-cell receptor beta-chain repertoires in the peripheral blood in response to concurrent chemoradiotherapy for Epstein-Barr virus-associated nasopharyngeal carcinoma. Oncoimmunology. 2021;10(1):1968172.

40. Wang G, Mudgal P, Wang L, et al. TCR repertoire characteristics predict clinical response to adoptive CTL therapy against nasopharyngeal carcinoma. Oncoimmunology. 2021;10(1):1955545.

41. Rosemarie Q, Sugden B. Epstein-Barr Virus: How Its Lytic Phase Contributes to Oncogenesis. Microorganisms. 2020;8(11).

42. Feng WH, Israel B, Raab-Traub N, Busson P, Kenney SC. Chemotherapy induces lytic EBV replication and confers ganciclovir susceptibility to EBV-positive epithelial cell tumors. Cancer Res. 2002;62(6):1920–1926.

43. Westphal EM, Blackstock W, Feng W, Israel B, Kenney SC. Activation of lytic Epstein-Barr virus (EBV) infection by radiation and sodium butyrate in vitro and in vivo: a potential method for treating EBV-positive malignancies. Cancer Res. 2000;60(20):5781–5788.

44. Grywalska E, Rolinski J. Epstein-Barr virus-associated lymphomas. Semin Oncol. 2015;42(2):291–303.

45. Ott PA, Dotti G, Yee C, Goff SL. An Update on Adoptive T-Cell Therapy and Neoantigen Vaccines. Am Soc Clin Oncol Educ Book. 2019;39:e70–e78.

46. Deng Y, Chatterjee B, Zens K, et al. CD27 is required for protective lytic EBV antigen-specific CD8+ T-cell expansion. Blood. 2021;137(23):3225–3236.

47. Tay RE, Richardson EK, Toh HC. Revisiting the role of CD4(+) T cells in cancer immunotherapy-new insights into old paradigms. Cancer Gene Ther. 2021;28(1-2):5-17.

48. Mautner J, Bornkamm GW. The role of virus-specific CD4+ T cells in the control of Epstein-Barr virus infection. Eur J Cell Biol. 2012;91(1):31–35.

49. Haque T, Wilkie GM, Jones MM, et al. Allogeneic cytotoxic T-cell therapy for EBV-positive posttransplantation lymphoproliferative disease: results of a phase 2 multicenter clinical trial. Blood. 2007;110(4):1123–1131.

50. Volk A, Hartmann S, Muik A, et al. Comparison of three humanized mouse models for adoptive T cell transfer. J Gene Med. 2012;14(8):540–548.

51. Rauser G, Einsele H, Sinzger C, et al. Rapid generation of combined CMV-specific CD4+ and CD8+ T-cell lines for adoptive transfer into recipients of allogeneic stem cell transplants. Blood. 2004;103(9):3565–3572.

52. Merlo A, Turrini R, Bobisse S, et al. Virus-specific cytotoxic CD4+ T cells for the treatment of EBV-related tumors. J Immunol. 2010;184(10):5895–5902.

53. Nowakowska J, Stuehler C, Egli A, et al. T cells specific for different latent and lytic viral proteins efficiently control Epstein-Barr virus-transformed B cells. Cytotherapy. 2015;17(9):1280–1291.

54. Khanna N, Stuehler C, Conrad B, et al. Generation of a multipathogen-specific T-cell product for adoptive immunotherapy based on activation-dependent expression of CD154. Blood. 2011;118(4):1121–1131.

