## Supplemental_materials for "Stem cell memory EBV-specific T cells control post-transplant lymphoproliferative disease and persist *in vivo*"

Supplementary methods

Supplementary figures 1-7

### SUPPLEMENTAL METHODS

#### Media used

All media contained PenStrep (Gibco) and were sterile-filtered when supplemented with the human serum (HS).

*CTL-M*

RPMI 1640 (Sigma Aldrich) with PenStrep (Gibco) and 5% HS.

*CTL-M with elevated $K^+$ concentration*

Powdered NaCl-free RPMI 1640 media (Gibco) was reconstituted as previously described[1]. Briefly, NaCl 63.4 mM, NaHCO3 23.8 mM, and *additional* KCl 40 mM had to be added for complete reconstitution. Then PenStrep and 5% HS was added.

*LCM-10*

RPMI 1640 (Gibco) with PenStrep 100x (Gibco), 5% human serum, 50x Glutamax (Gibco), 100x MEMNEA (Gibco) and 100x Sodium Pyruvate (Gibco).

*Concentrations of peptides, cytokines and TWS-119:*

| Reagent | Supplier | Final concentration |
|---|---|---|
| EBV Consensus pepmix | Miltenyi Biotec | 0.1 µg/peptide/mL |
| IL-2 | Proleukin | 20 U/mL |
| IL-4 | R&D | 400 U/mL |
| IL-7 | R&D | 10 ng/mL |
| IL-15 | Peprotech | 10 ng/mL |
| IL-21 | Peprotech | 30 ng/mL |
| TWS-119 | Selleckchem | 5µM |

*Flow cytometry and FACS panels*

1. General surface marker analysis (Cytek Aurora)

| **Dilution** | **Marker** | **Dye** | **Clone** | **Supplier** | **Catalog number** |
|---|---|---|---|---|---|
| Viability staining | | | | | |
| 1:1000 | Zombie Aqua | | | Biolegend | 423102 |
| Surface staining | | | | | |
| 1:400 | CD3 | BUV395 | UCHT1 | BD Biosciences | 563546 |
| 1:400 | CD4 | BUV496 | SK3 | BD Biosciences | 612936 |

| | | | | | |
|---|---|---|---|---|---|
| 1:500 | CD8 | BUV805 | SK1 | BD Biosciences | 612889 |
| 1:50 | CD19 | PE/Cy7 | HIB19 | Biolegend | 302216 |
| 1:30 | CD27 | FITC | M-T271 | Biolegend | 356403 |
| 1:100 | CD45 (anti-mouse) – only for in vivo samples | Alexa Fluor 647 | S18009D | Biolegend | 160304 |
| 1:50 | CD45 – only for in vivo samples | Pacific Blue | HI30 | Biolegend | 304029 |
| 1:50 | CD45RA | APC | MEM-56 | Thermo Fisher | MHCD45RA05 |
| 1:100 | CD45RO | Alexa Fluor 700 | UCHL1 | Biolegendd | 304218 |
| 1:30 | CD62L | BV650 | DREG-56 | Biolegend | 304832 |
| 1:33 | CD127 | BV605 | A019D5 | Biolegend | 351334 |
| 1:50 | CCR7 | APC/Cy7 | G043H7 | Biolegend | 353212 |
| 1:50 | CTLA4 | BV785 | BNI3 | Biolegend | 369624 |
| 1:50 | LAG3 | BV711 | 11C3C65 | Biolegend | 369320 |
| 0:200 | PD1 | BB700 | EH12.1 | BD Biosciences | 566460 |
| 1:100 | TIGIT | BV421 | A15153G | Biolegend | 372710 |
| 0:200 | TIM3 | BV480 | 7D3 | BD Biosciences | 746771 |

| | | | | | |
|---|---|---|---|---|---|
| 1:25 | MHC class I multimers | PE | - | MBL | See below |

2. Staining FACS-based sorting (BD FACSMelody)

| **Dilution** | **Marker** | **Dye** | **Clone** | **Supplier** | **Catalog number** |
|---|---|---|---|---|---|
| Surface staining | | | | | |
| 1:20 | CD3 | BV510 | UCHT1 | Biolegend | 300447 |
| 1:20 | CD27 | BV421 | M-T271 | Biolegend | 356418 |
| 1:20 | CD45RA | PE/ Dazzle 594 | HI100 | Biolegend | 356418 |
| 1:20 | CD45RO | FITC | UCHL1 | Biolegend | 304204 |
| 1:20 | CD62L | PE/Cy7 | DREG-56 | Biolegend | 304822 |

3. Flow cytometry after 7-day co-culture of sorted memory populations (Cytek Aurora)

| **Dilution** | **Marker** | **Dye** | **Clone** | **Supplier** | **Catalog number** |
|---|---|---|---|---|---|
| Whole cell staining (before co-culture start) | | | | | |
| 1:1000 | CellTrace Violet | | | Thermo Fisher | C34557 |
| Viability staining | | | | | |
| 1:1000 | Zombie UV | | | Biolegend | 423108 |
| Surface staining | | | | | |
| 1:20 | CD3 | BV510 | UCHT1 | Biolegend | 300447 |
| 1:20 | CD27 | BV421 | M-T271 | Biolegend | 356418 |

| | | | | | |
|---|---|---|---|---|---|
| 1:20 | CD45RA | PE/ Dazzle 594 | HI100 | Biolegend | 356418 |
| 1:20 | CD45RO | FITC | UCHL1 | Biolegend | 304204 |
| 1:20 | CD62L | PE/Cy7 | DREG-56 | Biolegend | 304822 |
| 1:25 | MHC class I multimers | PE | - | MBL | See below |
| Intracellular staining | | | | | |
| 1:50 | Granzyme B | PE/Cy5 | QA16A02 | Biolegend | 372226 |
| 1:50 | IFNγ | APC | 4S.B3 | Biolegend | 502512 |

4. Cytotoxicity assay (Cytek Aurora)

| **Dilution** | **Marker** | **Dye** | **Clone** | **Supplier** | **Catalog number** |
|---|---|---|---|---|---|
| Apoptosis staining (skipped for outgrowth assay) | | | | | |
| 1:1000 | CellEvent Caspase-3/7 Green Detection Reagent | | | Thermo Fisher | C10423 |
| Viability staining | | | | | |
| 1:1000 | Zombie Aqua | | | Biolegend | 423102 |
| Surface staining | | | | | |
| 1:50 | CD3 | APC | UCHT1 | Biolegend | 300412 |
| 1:50 | CD19 | PE/Cy7 | HIB19 | Biolegend | 302216 |

5. Weekly bleedings (BD LSRFortessa)

| **Dilution** | **Marker** | **Dye** | **Clone** | **Supplier** | **Catalog number** |
|---|---|---|---|---|---|
| Viability staining | | | | | |

| | | | | | |
|---|---|---|---|---|---|
| 1:1000 | Zombie Aqua | | | Biolegend | 423101 |
| Surface staining | | | | | |
| 1:400 | CD3 | PerCP/Cyanine5.5 | UCHT1 | Biolegend | 300429 |
| 1:400 | CD4 | APC | OKT4 | Biolegend | 317416 |
| 1:500 | CD8 | BV650 | SK1 | Biolegend | 344730 |
| 1:30 | CD19 | PE/Cy7 | HIB19 | Biolegend | 302216 |
| 1:50 | CD45 | Pacific Blue | HI30 | Biolegend | 304029 |
| 1:50 | CD45RA | BV785 | HI100 | Biolegend | 304139 |
| 1:30 | CD62L | FITC | DREG-56 | BD Biosciences | 555543 |
| 1:33 | CD127 | APC/Cy7 | A019D5 | Biolegend | 351348 |
| 1:50 | HLA-DR | BV605 | G46-6 | BD Biosciences | 562845 |
| 1:50 | PD-1 | PE/Dazzle 594 | EH12.2H7 | Biolegend | 329940 |
| Intracellular ad intranuclear staining | | | | | |
| 1:50 | Granzyme B | Alexa Fluor 700 | GB11 | BD Biosciences | 560213 |
| 1:50 | TCF-1 | PE | 7F11A10 | Biolegend | 655207 |

EBV-specific MHC class I multimer list

| Type | HLA Restriction | Peptide (PE-labeled) | Catalog number |
|---|---|---|---|
| Tetramer | HLA-A*02:01 | LMP2 356-364 FLYALALLL | TS-M069-1 |
| Tetramer | HLA-A*11:01 | LMP2 SSCSSCPLSK | TS-M111-1 |
| Tetramer | HLA-A*11:01 | EBNA3B 416-424 | TS-M029-1 |

| | | | |
|---|---|---|---|
| | | IVTDFSVIK | |
| Tetramer | HLA-A*02:01 | LMP2 426-434<br>CLGGLLTMV | TB-M032-1 |
| Tetramer | HLA-A*03:01 | EBNA3A 603-611<br>RLRAEAQVK | TB-M033-1 |
| Dextramer | HLA-A*02:01 | BMLF-1<br>GLCTLVAML | WB2130 |
| Dextramer | HLA-B*3501 | EBNA-1<br>HPVGEADYFEY | WK2145 |
| Dextramer | HLA-B*0702 | EBNA-3<br>RPPIFIRRL | WH2166 |
| Dextramer | HLA-B*0801 | EBNA-3A<br>FLRGRAYGL | WI2147 |

### Supplemental figures

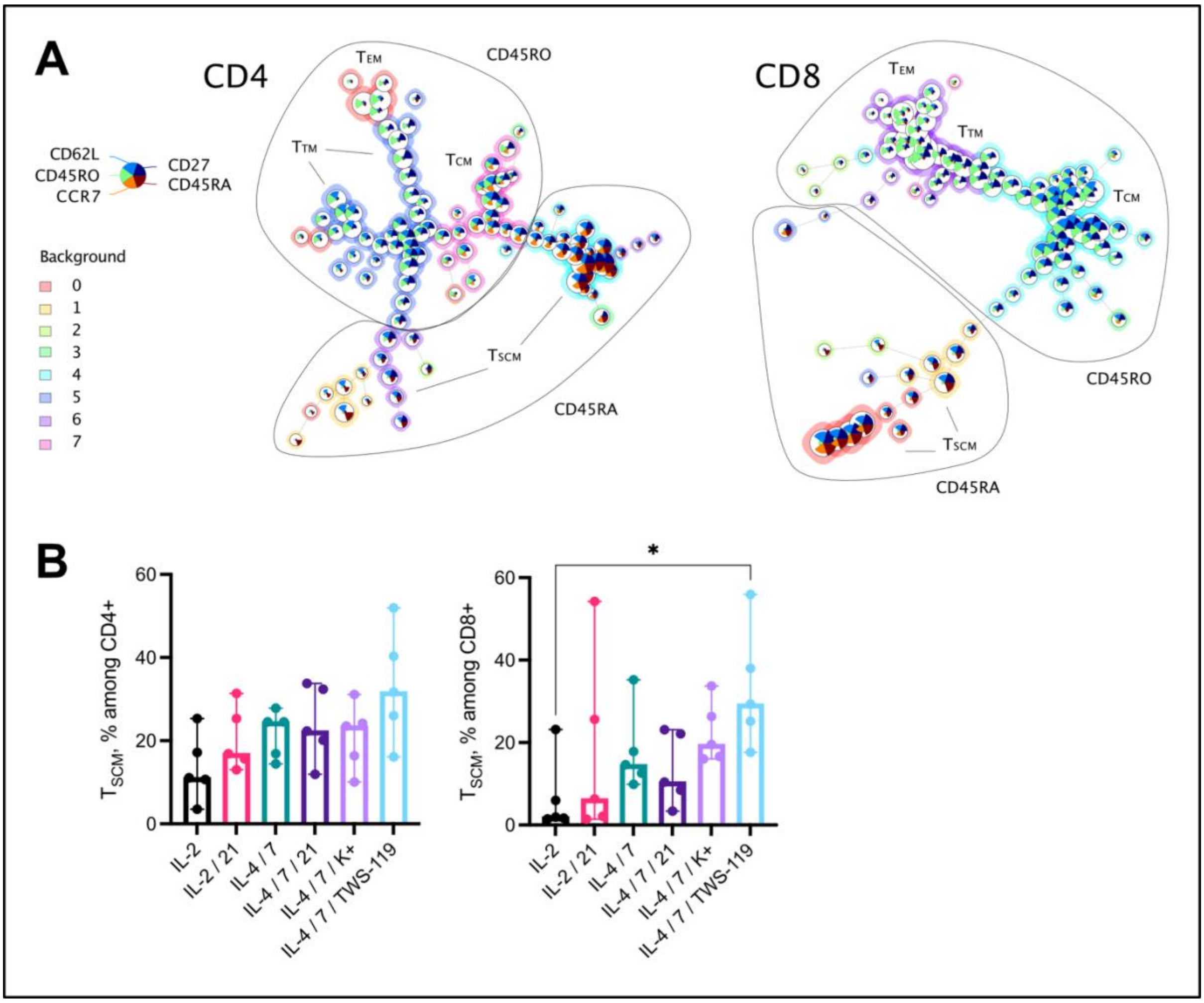


**Supplemental Figure 1. Defining markers for T cell memory phenotyping.** (A) Memory populations were distinguished among $CD4^+$ and $CD8^+$ T cells (separate algorithms for both) by CD27, CD45RA, CD45RO, CD62L and CCR7 markers in the FlowSOM algorithm. Based on the constructed maps, we could identify stem cell memory $T_{SCM}$ ($CD45RA^+CD27^+CD62L^+$), central memory $T_{CM}$ ($CD45RO^+CD27^+CD62L^+$), transitional memory $T_{TM}$ ($CD45RO^+CD27^+CD62L^-$) and effector memory $T_{EM}$ ($CD45RO^+CD27^-CD62L^-$). Short-lived terminally differentiated TEMRA populations ware almost negligible and, in the analysis, they were defined as ($CD45RA^+CD27^-CD62L^-$). CCR7 was less expressed than CD62L among $CD8^+$ T cells and thus was excluded for simplicity. (B) Stem cell memory T cell proportions among expanded $CD4^+$ and $CD8^+$ T cells; n=5, medians with range, Friedman test, $\alpha=0.05$, $*p < 0.05$.

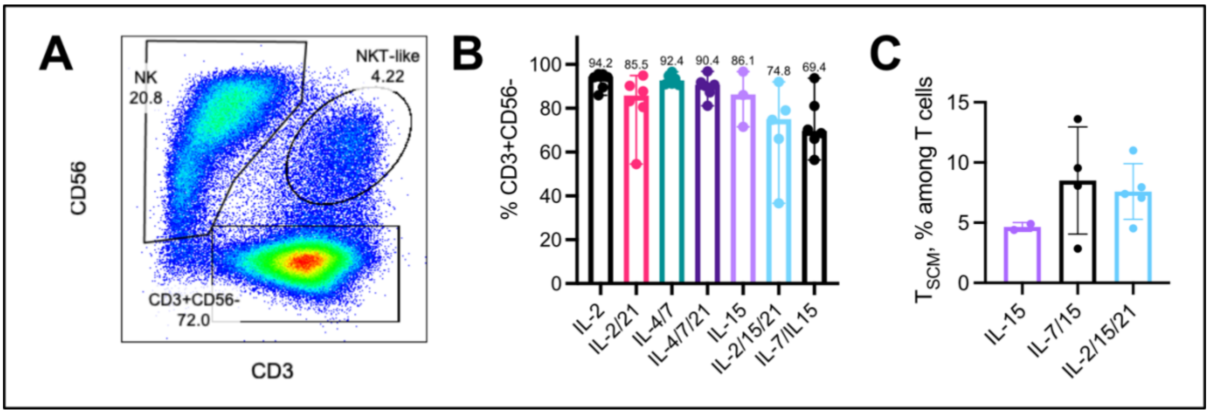


**Supplemental Figure 2. NK cell enrichment during PBMC culture in the presence of IL-15.** (A) Representative plot showing enrichment of CD56$^+$CD3$^-$ NK cells in an EBV-CTL product. (B) The purity (percentage of CD3$^+$CD56$^-$ among live cells) of resulting products. (C) Proportions of $T_{SCM}$ populations.

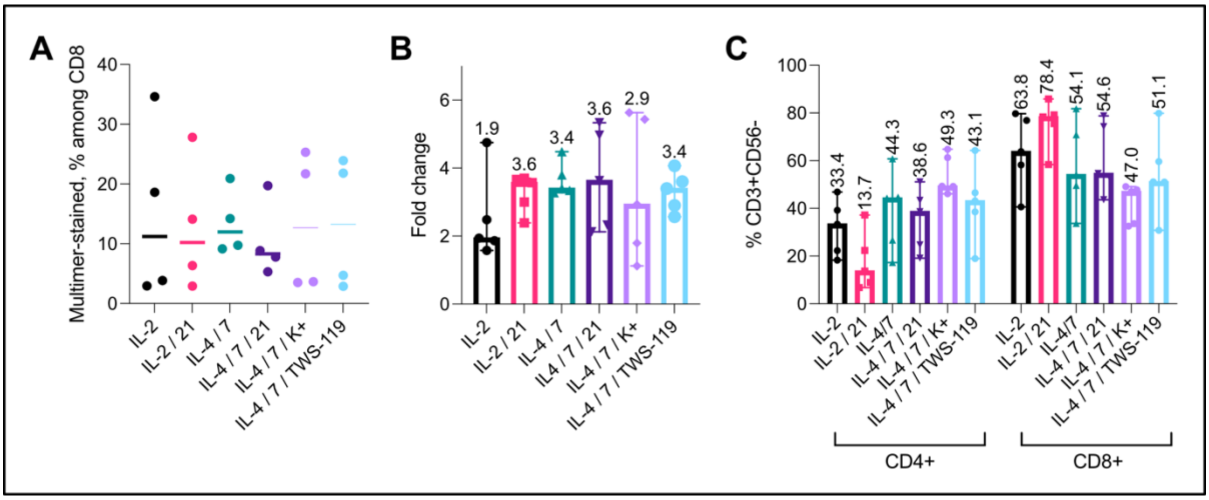


**Supplemental Figure 3. General characteristics of EBV-CTL products.** (A) Proportions of MHC class I-peptide specific CD8$^+$ T cells measured by respective multimer stainings. N=4, medians. (B) Total cell expansion in fold change, and (C) CD4$^+$ and CD8$^+$ T cell proportions of EBV-CTL products; n=5, medians with range.

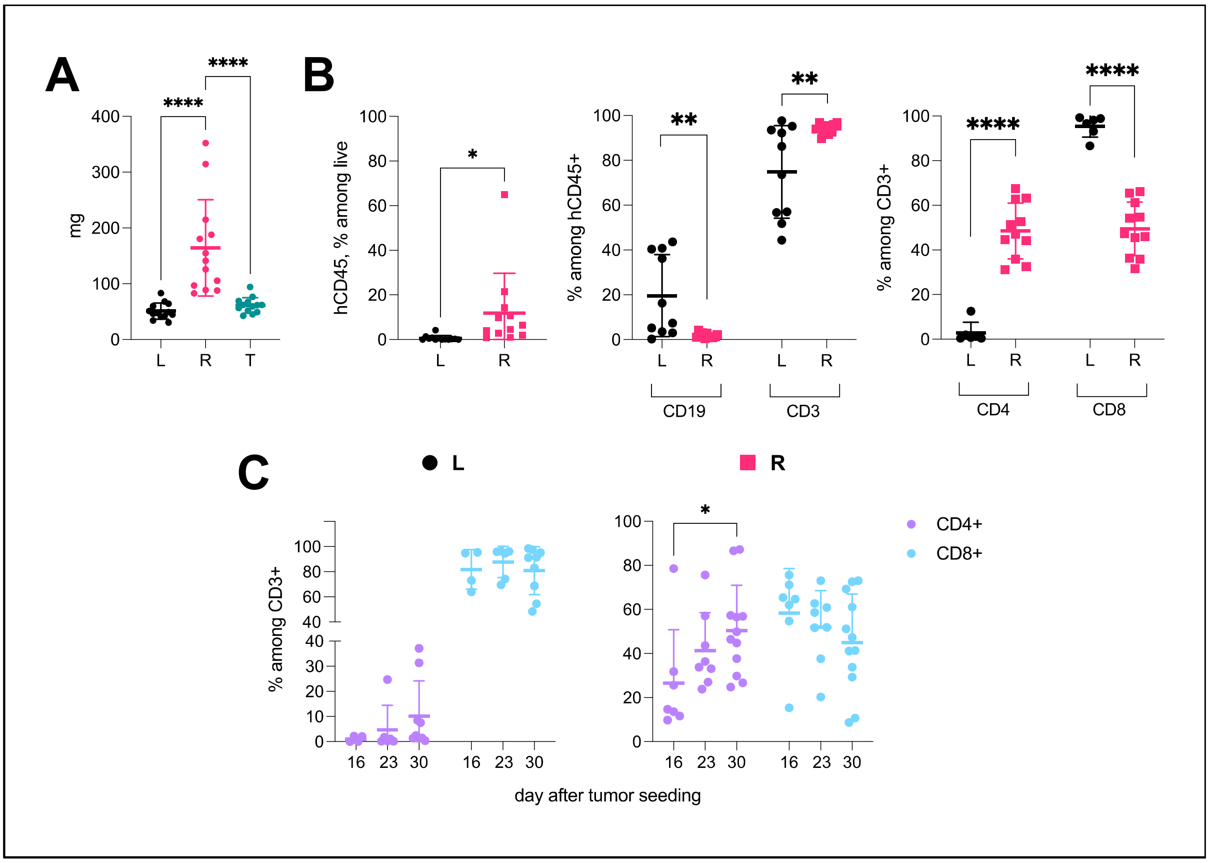


**Supplemental Figure 4. General characteristics of *in vivo* experiments.** (A) spleen weights. (B) human CD45$^{+}$, CD19$^{+}$ vs CD3$^{+}$, CD4$^{+}$ vs. CD8$^{+}$ cell proportions in bone marrow. (C) Dynamic of CD4$^{+}$ and CD8$^{+}$ T cell frequency changes over the experimental time course in peripheral blood. For A-C, mixed effects analysis was used.

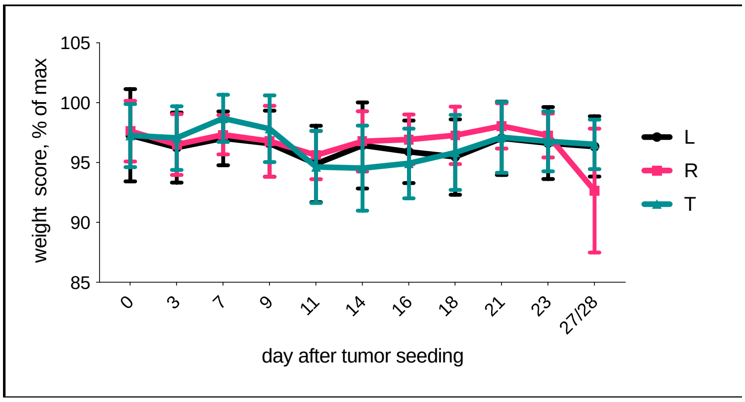


**Supplemental Figure 5. Weight loss in the CTL-R and control groups and associated parameters.** Weight score dynamics during in vivo experimentations, shown are means with SD, mixed effects analysis (no significance).

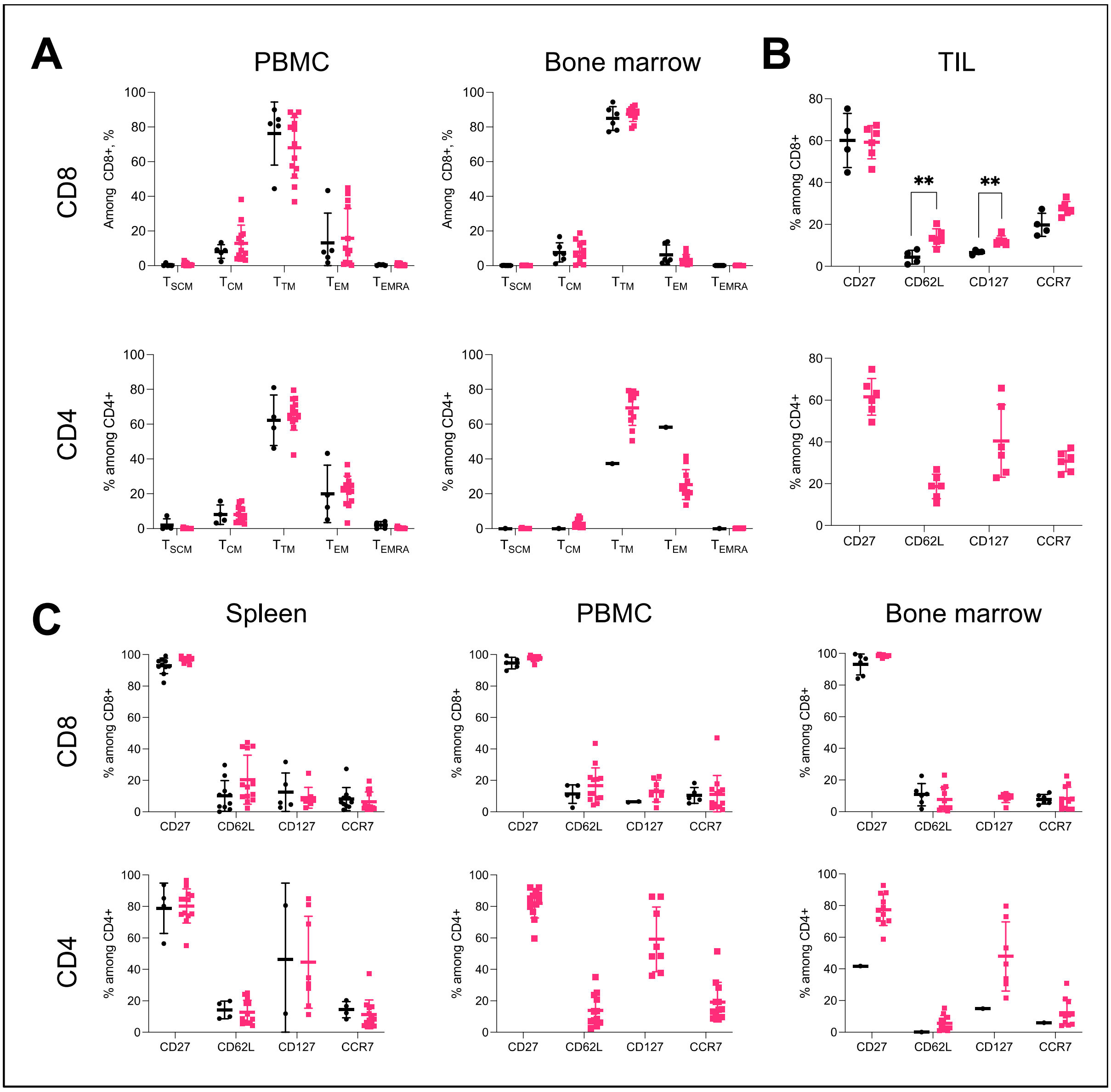


**Supplemental Figure 6. Memory T cell phenotyping in different organs from *in vivo* experiments.** CTL-L groups are shown in black circles, and CTL-R – in pink squares. Only samples with >100 events / population are shown. (A) Memory T cell populations in peripheral blood (PBMCs) and bone marrow. Expression of separate memory markers in TILs (B) and spleen, PBMCs and bone marrow (C).

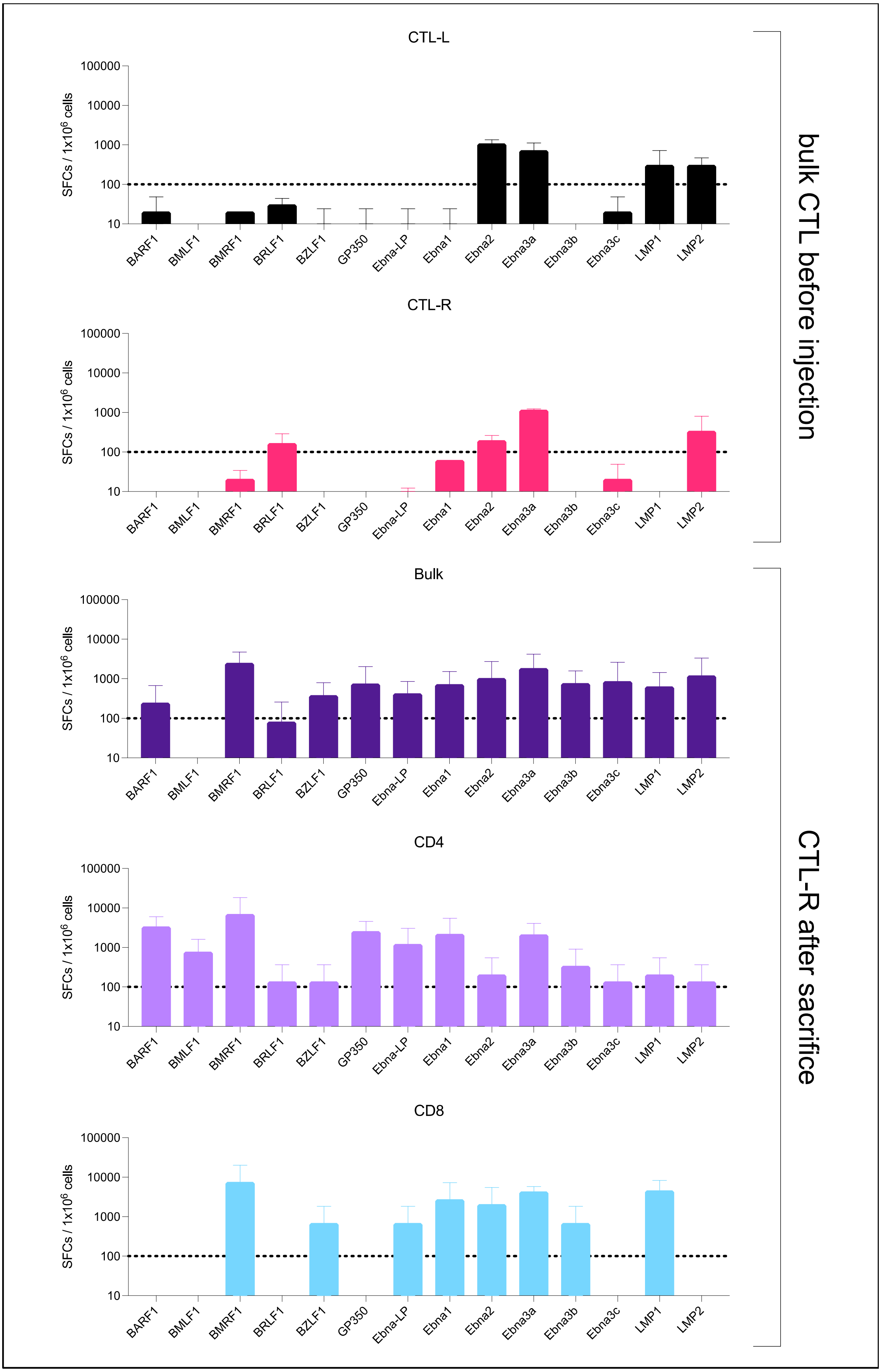
CTL-L
CTL-R
Bulk
CD4
CD8
SFCs / 1x10^6 cells
100000
10000
1000
100
10
BARF1
BMLF1
BMRF1
BRLF1
BZLF1
GP350
Ebna-LP
Ebna1
Ebna2
Ebna3a
Ebna3b
Ebna3c
LMP1
LMP2
bulk CTL before injection
CTL-R after sacrifice

98

**Supplemental Figure 7. Cellular IFNγ secretion in infusion product and after adoptive transfer in CTL-R group.** Quantitative details of EBV specific responses of bulk CTL-L and CTL-R cultures before injection vs CTL-R bulk, $CD4^+$ and $CD8^+$ splenocytes after sacrifice in response to stimulation with single-EBV protein derived peptide pools and measured by IFNγ ELISpot assays (based on mean data from all available samples collected from one experiment / one donor).


1. Vodnala, S.K. et al. T cell stemness and dysfunction in tumors are triggered by a common mechanism. *Science* **363** (2019).
2. Nowakowska, J. et al. T cells specific for different latent and lytic viral proteins efficiently control Epstein-Barr virus-transformed B cells. *Cytotherapy* **17**, 1280-1291 (2015).
3. Khanna, N. et al. Generation of a multipathogen-specific T-cell product for adoptive immunotherapy based on activation-dependent expression of CD154. *Blood* **118**, 1121-1131 (2011).
